# The microRNA regulatory landscape of skeletal muscle-derived extracellular vesicles

**DOI:** 10.1101/2025.06.02.657546

**Authors:** Yudai Kawamoto, Atomu Yamaguchi, Xiaoqi Ma, Hidemi Fujino, Noriaki Maeshige

## Abstract

Skeletal muscle-derived extracellular vesicles (SkM-EVs) have recently been recognized as novel endocrine factors capable of facilitating inter-organ communication between skeletal muscle and distant organs. These vesicles transport various molecular cargoes, including microRNAs (miRNAs), which are essential regulators of post-transcriptional gene expression. In this study, we characterized the miRNA composition of SkM-EVs using small RNA sequencing and elucidated their putative biological roles via comprehensive bioinformatics analyses. Gene Ontology (GO) and Kyoto Encyclopedia of Genes and Genomes (KEGG) pathway enrichment analyses of the predicted miRNA targets revealed that SkM-EV miRNAs are involved in several key pathways, including the FoxO signaling pathway, the insulin signaling pathway, and cancer-related pathways. Our findings suggest that SkM-EV miRNAs may simultaneously promote muscle differentiation and exert protective effects against diabetes and cancer development. These findings provide new insights into the systemic regulatory roles of SkM-EVs and highlight their therapeutic potential for muscular, metabolic, and oncological disorders.

## 1. Introduction

Skeletal muscle, traditionally regarded solely as a locomotor organ, has emerged as the largest endocrine tissue, secreting myokines and exerkines (1–3). This organ significantly contributes to systemic metabolism, accounting for approximately 75% of whole-body metabolic activity (4). In particular, skeletal muscle-derived extracellular vesicles (SkM-EVs) have recently gained attention as endocrine mediators secreted by skeletal muscle (5). EVs are lipid-bilayer vesicles that carry proteins, lipids, nucleic acids, mRNA, and microRNAs (miRNA) from donor cells. These cargoes facilitate intercellular communication (6).

Skeletal muscle lies on the surface of the body and can be contracted voluntarily, enabling easy, noninvasive stimulation via exercise (7), electrical stimulation (8), or ultrasound (9). Indeed, it has been reported that high-intensity exercise with muscle contraction increases the amount of circulating EVs (10) and that high-intensity ultrasound stimulation to cultured myotubes promotes EV secretion (11). Thus, the release of skeletal muscle– derived EVs can be intentionally and precisely regulated by selecting the appropriate stimulation modality. Since SkM-EV production can be precisely enhanced by selecting the appropriate stimulation, SkM-EVs are promising candidates for therapeutic application.

miRNAs regulate the expression of approximately 80% of human genes through base-pairing of their “seed” sequences with complementary mRNAs (12). After EVs are internalized into target cells or organs, miRNAs can regulate gene expression and promote physiological effects (13–15). This miRNA cargo makes up an essential part of EV profiles and is key contributor to the overall biological function of EVs. However, SkM-EVs carry hundreds of miRNAs, making it difficult to attribute specific phenotypic outcomes to individual miRNAs. Moreover, cells adjust both the release dynamics and molecular content of EVs in response to the cellular microenvironment (16); for example, muscle injury triggers selective miRNA packaging rather than passive leakage (17). To understand EVs from dysfunctional skeletal muscle, we first need a comprehensive profile of miRNAs in EVs released by healthy skeletal muscle. Previous studies mostly used a candidate approach—focusing on single miRNAs with reported therapeutic effects: anti-inflammatory effects (18), muscle regeneration (14), bone remodeling (19,20), angiogenesis (21), or proliferation (22). However, this strategy may underestimate broader functions of EVs on whole-body homeostasis.

In this study, we quantified SkM-EV miRNAs and assessed their impact on whole-body metabolism via bioinformatics analysis (Figure 1).

**Figure 1.**
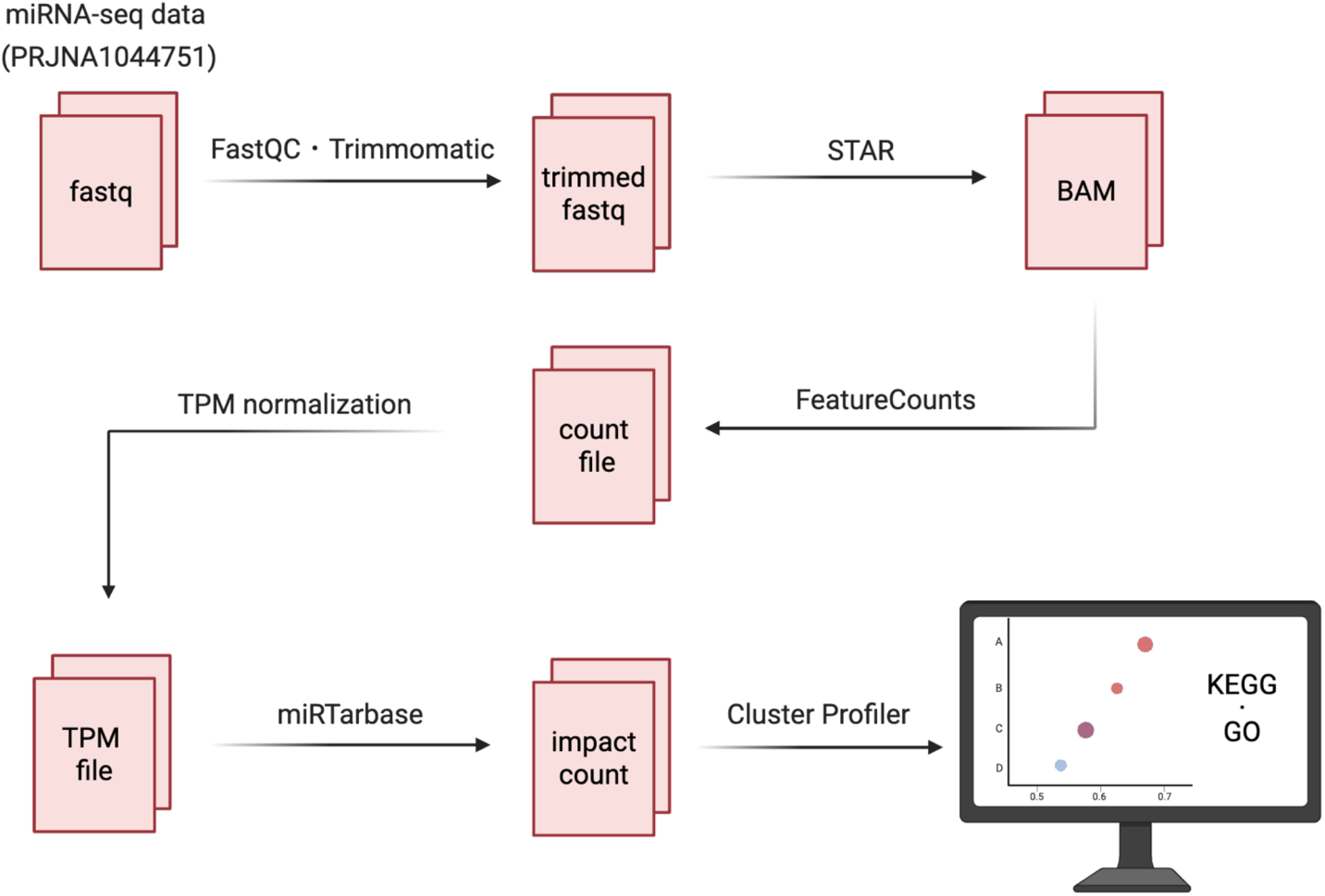
Overview of miRNA-Seq data analysis pipeline for PRJNA1044751. Red boxes den2o6t7e the output files generated at each step, and labels above the arrows indicate the software tools268 used or the processing steps performed.

## 2. Materials and methods

### 2.1. Data acquisition

We analyzed miRNA sequencing (miRNA-seq) data from the previously generated single-end RNA libraries prepared from SkM-EV samples of the C2C12 cell line (BioProject PRJNA1044751). Briefly, miRNAs were isolated from myotube-derived EVs using TRIzol reagent (Takara Biotechnology, Japan) according to the manufacturer’s protocol. Libraries were sequenced on an Illumina NovaSeq 6000 platform.

### 2.2. Bioinformatics analysis of miRNA-seq data

miRNA-seq data quality was assessed using FastQC (v0.12.1). Trimmomatic (v0.39) (23) was used to trim sequencing adapters (AGATCGGAAGAGCACACGTCT) from the 3’-end. Reads were aligned to the mm10 mouse genome using STAR (v2.7.11b) (24), with microRNA annotations from miRBase (v22.1). FeatureCounts (25) was used to analyze the resulting BAM files.

### 2.3. Identification of miRNA targets

Experimentally validated miRNA–mRNA interactions were retrieved from miRTarBase (26). SkM-EV miRNAs were quantified as transcripts per million (TPM). miRNAs with TPM < 10 were excluded. The subsequent conversion to mRNA was accompanied by the calculation of impact counts, which were derived as the cumulative TPM of corresponding miRNAs.

### 2.4. Pathway analyses of miRNA targets

Targeted mRNAs were ranked by converted impact counts and analyzed by Gene set enrichment analysis (GSEA). Simultaneously, impact counts of mRNA data underwent a log2(TPM + 1) transformation to stabilize calculations. Gene Ontology (GO) analysis was performed to classify genes into cellular components, molecular functions, and biological processes, while Kyoto Encyclopedia of Gene and Genomes (KEGG) pathway enrichment analysis evaluated the functional roles of genes (27,28). Both analyses were conducted using the Bioconductor package ClusterProfiler (29).

Benjamini-Hochberg corrected p-values were calculated for KEGG and GO analyses using 1,000 permutation tests. Pathways with *adjusted p-value* < 0.05 were considered significantly enriched.

### 2.5. Statistical analysis

ggplot2 (30) was used to generate bar graphs with average and standard deviation (SD) of repeated experiments. *p* < 0.05 was considered statistically significant.

## 3. Results

### SkM-EV miRNAs are involved in diverse biological processes

As previously demonstrated (29), SkM-EV samples comprised 745 distinct miRNA species spanning a broad dynamic range of abundance (Figure 2A). Experimentally validated miRNA–mRNA interactions were retrieved from miRTarBase (Figure 2B). For each miRNA detected in SkM-EVs, we identified its target mRNAs (n = 11,887), quantified their expression in TPM, and then converted those TPM values into impact counts for downstream analysis. A detailed breakdown is provided in Supplementary File.

**Figure 2.**
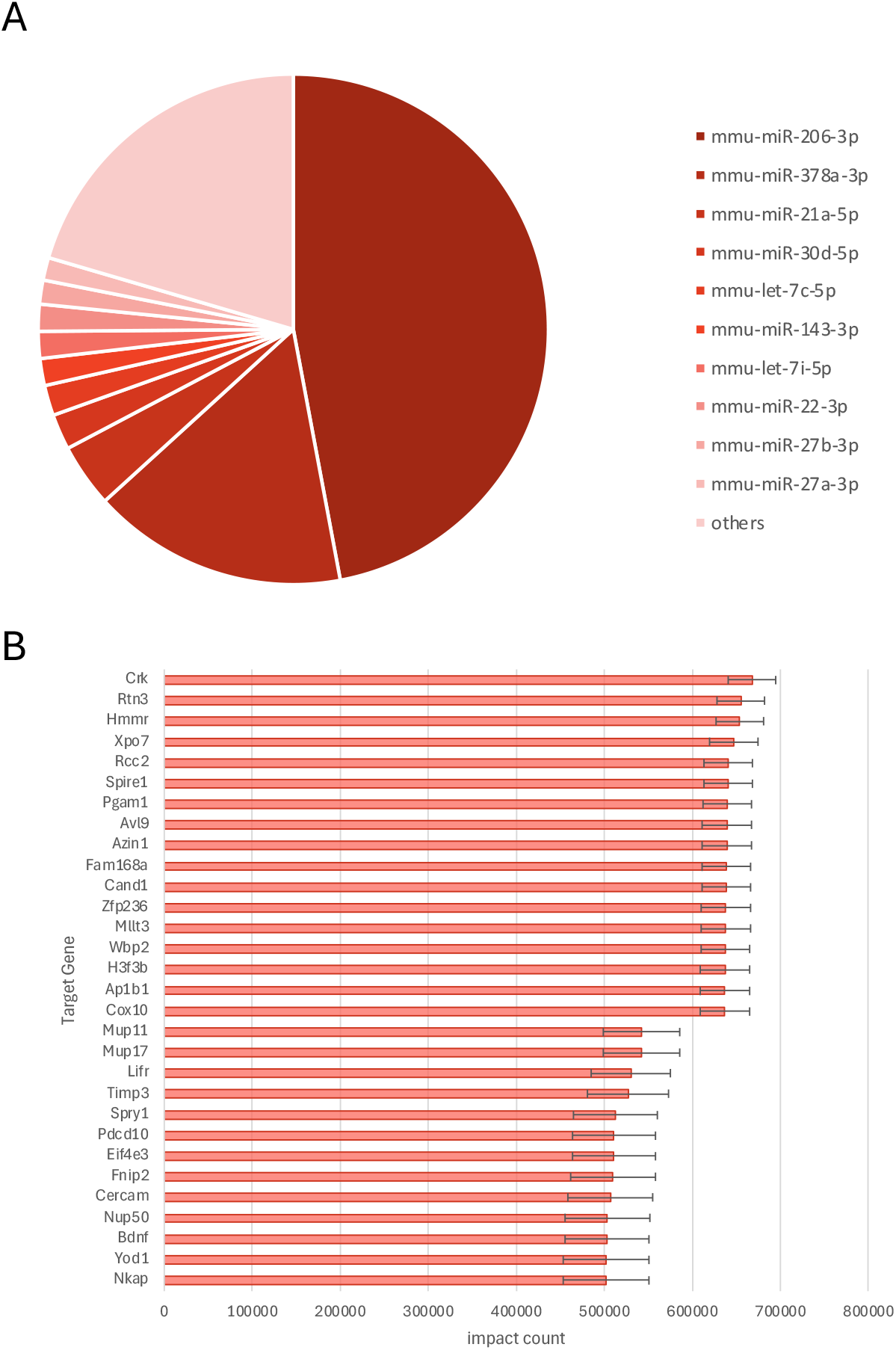
**m**iRNA expression in SkM-EVs; (A) The two most abundant miRNAs accounted for 63.7% of the total miRNA profile (n = 3). (B) Read distribution of top 30 target genes.

KEGG GSEA analysis demonstrated that target genes are significantly implicated in a multitude of signaling pathways, including Neurotrophin, Insulin, Relaxin, FoxO, AGE-RAGE and PI3K-Akt; additionally, these genes are involved in the regulation of various cancer-related pathways, such as small cell lung cancer and proteoglycans in cancer (Figure 3A, B). GO enrichment analysis provided further insights (Figure 3C), revealing that the target genes are primarily associated with the following biological processes: “negative regulation of cardiocyte differentiation,” “extrinsic apoptotic signaling pathway via death domain receptors,” “insulin receptor signaling pathway,” “cellular response to insulin stimulus,” and “target genes for the insulin receptor.” In terms of the cellular component, “RSC-type complex,” “complex of collagen trimers,” and “dendritic spine” were enriched. Additionally, molecular functions such as “mRNA binding” and “protease binding” were also found to be enriched.

**Figure 3.**
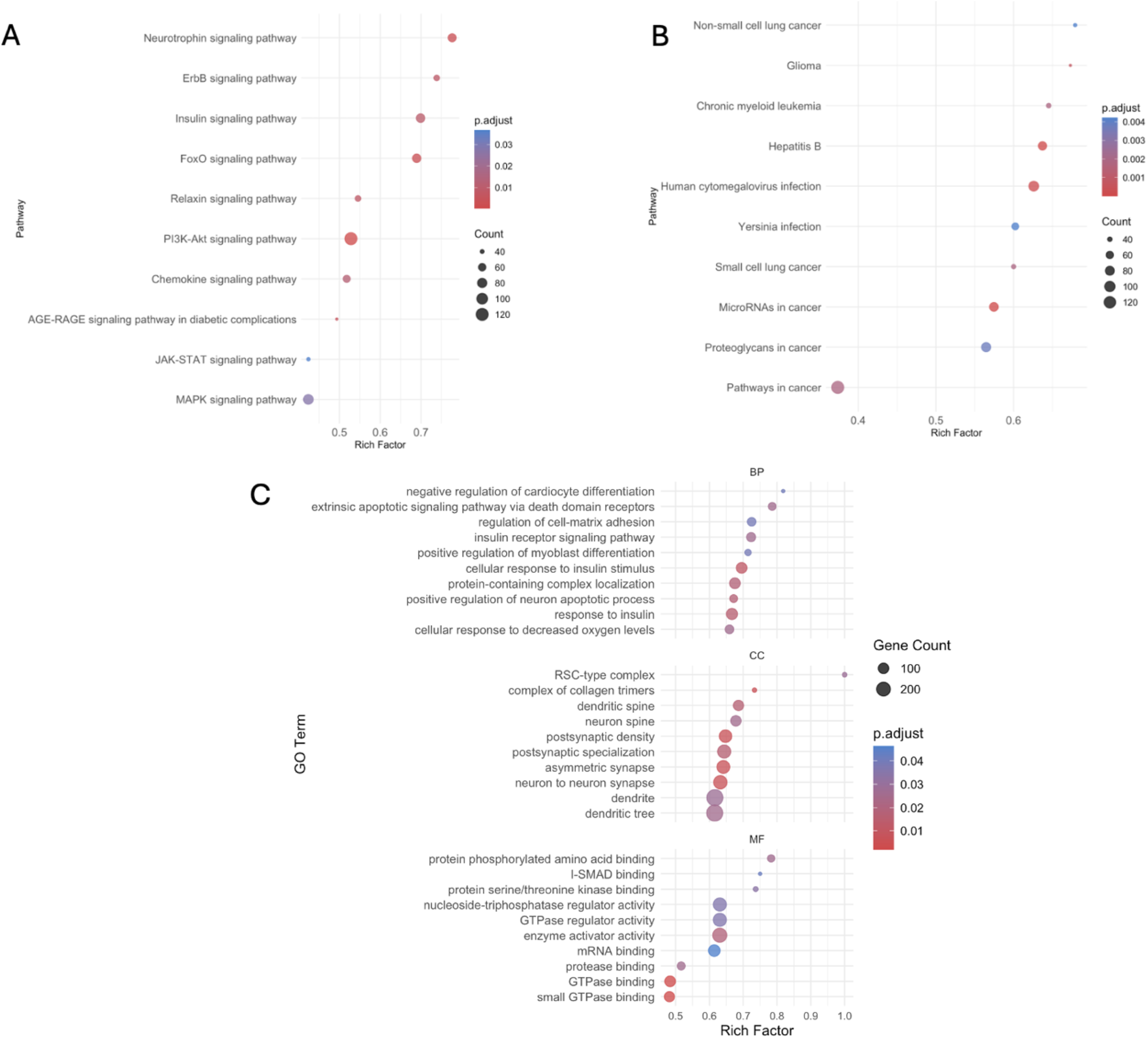
KEGG and GO enrichment analysis: (A) KEGG signaling, (B) KEGG non-signaling pathways, and (C) GO categories of miRNA targets, using GSEA.

## 4. Discussion

### 4.1. General overview

Skeletal muscle-derived extracellular vesicles (SkM-EVs) have demonstrated numerous beneficial properties,including anti-inflammatory effect (18), muscle regeneration (14), bone remodeling (19,20), promotion of angiogenesis (21) and promotion of proliferation (22). SkM-EVs are anticipated to have diverse therapeutic applications. However, these validated effects are limited, because previous studies have employed a candidate approach focusing on specific miRNAs to assess their therapeutic potential. In this study, we employed a bioinformatics approach to comprehensively elucidate the diverse biological effects exerted by miRNAs contained within SkM-EVs, considering their expression levels for the first time.

The results suggest that SkM-EVs possess a multifaceted range of functions.

### 4.2. SkM-EV miRNAs promote regeneration and differentiation of skeletal muscle

Skeletal muscles are responsible for physical activities, including excercise and life activities. In addition, skeletal muscle participates in the uptake, utilization, and storage of metabolic substrates such as glucose, lipids, and amino acids, accounting for up to 75% of whole-body metabolism (4). However, inactivity, aging, and hereditary factors can cause an imbalance in the synthesis and degradation of skeletal muscle proteins, leading to muscle wasting and eventual atrophy. Atrophy negatively affects daily life by impairing skeletal muscle metabolism, which reduces the ability to perform daily activities and prolongs recovery time from illness (31). GO analysis of BPs identified “regulation of extracellular matrix adhesion.” KEGG analysis revealed that miRNA targets significantly suppress the “FoxO signaling pathway and “JAK-STAT signaling pathway.” Because FoxO3 activation via atrogin-1 upregulation (32) and JAK–STAT3 activation (33) both induce muscle atrophy, SkM-EVs may prevent atrophy, support cell growth, and promote regeneration. In terms of specific miRNAs, Forterre et al. revealed that miR-22, miR-181a, and miR-133a in SkM-EVs facilitate muscle differentiation by repressing the expression of Sirt1, a known inhibitor of myogenesis. (14). miR-22 also regulates the PTEN/AKT/FoxO1 signaling pathway, blocking FoxO1 activity and suppressing FoxO3-induced muscle atrophy (34). miR-206, which accounts for over 47% of miRNAs in SkM-EVs, promotes myogenic differentiation by suppressing inhibitors of muscle cell maturation—such as a subunit of DNA polymerase— and by downregulating follistatin-1 (Fstl1), utrophin (Utrn), and Pax7, thereby facilitating the conversion of fibroblasts into skeletal muscle cells (35,36). Furthermore, I-SMAD binding, as revealed by GO analysis of molecular functions (MF), promotes skeletal muscle formation by inhibiting the TGF-β signaling pathway (37). Despite low levels of miR-181a and miR-133a, our results indicate that SkM-EVs are key players in muscle regeneration and differentiation.

### 4.3. SkM-EV miRNAs Modulate Insulin Signaling and Glucose Homeostasis

Skeletal muscle is the primary insulin-dependent tissue for glucose uptake, playing a central role in systemic glucose and energy homeostasis (38). However, in conditions like diabetes and dyslipidemia, insulin resistance impairs muscle glucose uptake, leading to hyperglycemia and impaired glucose homeostasis (39).

We found that “Insulin signaling pathway” and “AGE-RAGE signaling pathway in diabetic complications” in KEGG analysis and “Response to insulin,” “Cellular response to insulin stimulus” and “Insulin receptor signaling pathway” in BP of GO analysis were significantly suppressed by miRNAs in SkM-EV. These results suggest that SkM-EVs are closely involved in insulin signaling and glucose metabolism. Khalilian et al. reported KEGG enrichment of miR-206 target genes demonstrated their involvement in the “AGE-RAGE signaling pathway in diabetic complications” (40). miR-206 promotes insulin signaling by inhibiting PTPN1 expression and plays a role in improving insulin resistance in diabetes (41). miR-378 maintains hepatic glucose and lipid homeostasis through regulation of the PI3K subunit p110α (42), activates pyruvate-PEP futile cycle via the miR-378-Akt1-FoxO1-PEPCK pathway in skeletal muscle, and induces lipolysis in adipose tissues mediated by miR-378-SCD1 (43). Moreover, miR-378 enhances insulin-mediated glucose uptake in skeletal muscle and maintains glucose homeostasis in liver and skeletal muscle(44). Notably, fluorescently labeled SkM-EVs injected intramuscularly accumulate in the liver, spleen, pancreas, muscle, kidney, and gastrointestinal tract (22), directly enhancing of insulin secretion and sensitivity in these organs. Thus, SkM-EVs may function as a novel, noninvasive, and continuous drug delivery system for diabetes treatment.

### 4.4. SkM-EV miRNAs inhibit cancer growth and progression

Cancer is a complex, multifactorial disease defined by the sequential acquisition of functional capabilities that drive normal cells toward malignant transformation. These capabilities include sustained proliferative signaling, evasion of growth suppressors, resistance to cell death, replicative immortality, induction of angiogenesis, activation of invasion and metastasis, metabolic reprogramming, and immune evasion (45). Moreover, cancer induces cachexia, a metabolic syndrome characterized by the sustained loss of skeletal muscle and adipose tissue, which severely impairs everyday activities and represents a critical determinant of patient quality of life (46).

We have identified pathways associated with cancer: “Non-small cell lung cancer,” “miRNAs in cancer,” “Pathways in cancer,” “glioma,” “Proteoglycans in cancer” and “Small cell lung cancer.” Chen et al. revealed miR-206 represses HGF-induced epithelial-mesenchymal transition and angiogenesis in non-small cell lung cancer via c-Met /PI3K/Akt/mTOR pathway (47), which was among the pathways significantly suppressed in our KEGG analysis. Most tumors activate the MAPK and PI3K/AKT pathways. MAPK signaling, in particular, regulates key steps in angiogenesis and inflammatory-factor induction—processes driven by PI3K/AKT activation during both early and late stages of tumorigenesis. (48). Furthermore, miR-378 inhibits colorectal, prostate, and liver tumor growth (49). These findings indicate that SkM-EVs may exert anti-tumor effects across multiple cancer types, beyond non–small cell lung cancer.

## 5. Conclusion

We have shown the potential efficacy of SkM-EVs in the treatment of muscle regeneration, diabetes, and cancer pathologies by bioinformatics analysis. Still, the current research does not provide sufficient evidence to confirm the effects of SkM-EVs on cancer and diabetes; consequently, further studies are necessary to elucidate these potential effects.

## Notes

https://1drv.ms/f/c/9a9189d362efa69e/EmU8aMlzK3NBt-GxbYyaKSEBcxTonT4YRMm4l9wN1l2HOw

### Competing Interest Statement

The authors have declared no competing interest.

